# Acute decrease in plasma membrane tension induces macropinocytosis *via* PLD2 activation

**DOI:** 10.1101/594416

**Authors:** Julie Loh, Jophin Joseph, Mei-Chun Chuang, Shan-Shan Lin, Yu-Chen Chang, You-An Su, Allen P. Liu, Ya-Wen Liu

## Abstract

Internalization of macromolecules and membrane into cells through endocytosis is critical for cellular growth, signaling, and membrane tension homeostasis. Although endocytosis is responsive to both biochemical and physical stimuli, how physical cues modulate endocytic pathways is less understood. In contrary to the accumulating discoveries on effects of increased membrane tension on endocytosis, little is known about how a drop of tension impacts membrane trafficking. Here we reveal that acute reduction of plasma membrane tension results in phosphatidic acid, F-actin and dynamin 2-enriched dorsal membrane ruffling and subsequent macropinocytosis in myoblast. The membrane flaccidity-induced local phosphatidic acid production depends on phospholipase D2 (PLD2) that is activated *via* lipid raft disruption. Furthermore, the “membrane flaccidity-PLD2-macropinocytosis” pathway is dominant in myotube, reflecting a potential mechanism of membrane tension homeostasis upon intensive muscle stretching and relaxation. Together, we identify a new mechanotransduction pathway which converts acute tension drop into PA production and subsequently initiates macropinocytosis *via* actin and dynamin activities.

**Summary:** We reveal a mechanical induction of macropinocytosis that is elicited by acute decrease of plasma membrane tension, followed by lipid raft destabilization, PLD2 activation and PA production.

## Introduction

Eukaryotic cells harness multiple endocytic pathways to internalize fluid and membrane into transport vesicles from the plasma membrane (Conner and Schmid, 2003; Doherty and McMahon, 2009). In response to various cellular demands and environmental stimuli, endocytic machineries are regulated biochemically or physically to govern the growth and survival of cells (Dai and Sheetz, 1995; Liu et al., 2017; Scita and Di Fiore, 2010). Among them, macropinocytosis is an endocytic pathway that is induced by biochemical stimuli, including nutrients, growth factors, integrin substrates or even viruses and forms actin-based membrane ruffling and engulfment that leads to the formation of macropinosomes (Buckley and King, 2017; Doherty and McMahon, 2009).

Upon hyper-stimulation of growth factor receptors, activated PI3K and small GTPases initiate macropinocytosis *via* phosphatidylinositol (3,4,5)-trisphosphate (PIP_3_) production and actin polymerization (Levin et al., 2015; Yoshida et al., 2018). After membrane ruffling and closure, a sealed and large endocytic vacuole (1-10 μm in diameter) is formed. With the relatively large amount of solutes and membrane area being internalized, macropinocytosis is an efficient and ideal route to quickly uptake nutrients or attenuate cell signaling (Doherty and McMahon, 2009). Despite the clearly-defined, biochemically induced mechanism of macropinocytosis, little is known about the effects of mechanical alteration on macropinosome formation.

Membrane tension, the force applied on plasma membrane, has emerged as a master integrator which governs distinct cellular processes, including membrane trafficking, cell migration, immunological responses, cell growth and differentiation (Dai and Sheetz, 1995; Diz-Munoz et al., 2013; Masters et al., 2013). For endocytosis, cells need to overcome higher energetic barrier to bend the membrane inwardly when membrane tension is high, *e.g.* by using the pulling force from actin polymerization in clathrin-mediated endocytosis. (Boulant et al., 2011; Tan et al., 2015; Weinberg and Drubin, 2012). Conversely, when tension is decreased, it is easier for endocytic machinery to deform the membrane thus endocytosis is increased (Saleem et al., 2015; Shi and Baumgart, 2015).

Plasma membrane tension is contributed by membrane-cytoskeleton adhesion and osmotic pressure (Gauthier et al., 2012). Although membrane tension is used to control multiple cellular events, cells also need to maintain the homeostasis of membrane tension by sensing the change of tension and modulating its membrane area or cytoskeletal attachment *via* endocytosis, exocytosis, membrane invagination, actin polymerization or depolymerization (Diz-Munoz et al., 2013; Gauthier et al., 2012; Nassoy and Lamaze, 2012). Together, a feedback regulation of membrane tension, actin polymerization and membrane trafficking underpins the tension homeostasis and membrane remodeling events in cells.

The sensors of membrane tension are generally transmembrane proteins or peripheral membrane proteins, including stretch-activated ion channels, caveolin-cavin complex, curvature sensing proteins or membrane-actin interacting proteins (Diz-Munoz et al., 2013; Tsujita et al., 2015). Recently, Petersen *et al*. reported that lipid raft could function as a mechanosensor that undergoes kinetic or mechanical disruption to activate phospholipase D2 (PLD2) (Petersen et al., 2016). PLD2 is a plasma membrane-localized lipase which catalyzes the conversion of phosphatidylcholine (PC) into phosphatidic acid (PA) and choline, and is involved in endocytosis and actin polymerization (Antonescu et al., 2010; Colley et al., 1997; Du et al., 2004; Jiang et al., 2016). PLD2 mainly localizes at a lipid microdomain, the lipid raft, on plasma membrane *via* palmitoylation on its C223 and C224 residues where PC and phosphatidyl-4,5-bisphosphate (PI(4,5)P_2_), its substrate and activator, are excluded (Xie et al., 2002). Chemical or mechanical disruption of lipid raft may release the segregated PLD2 thus enhance the PA production at the plasma membrane (Diz-Munoz et al., 2016; Petersen et al., 2016).

PA is a negatively charged, cone-shaped lipid identified as a key mediator in phospholipid metabolism, mTOR activation, membrane trafficking, mitochondrial fusion and actin polymerization (Liu et al., 2013; Yang and Frohman, 2012). PA could be produced through phospholipid synthesis (the Kennedy pathway) in the endoplasmic reticulum, the phosphorylation of diacylglycerol by diacylglycerol kinase, or the hydrolysis of PC by PLD on plasma or endosomal membranes (Liu et al., 2013). Given the activity of PLD2 regulated by mechanical cues as well as the importance of its enzymatic product, PA, on actin polymerization and membrane trafficking, we hypothesize that alteration of membrane tension may affect PLD2 activity thus impacts endocytosis. Here, we discover that an acute decrease of plasma membrane tension induces PA production which triggers macropinocytosis without eliciting an increase of PIP_3_.

## Results

### Enrichment of PA at dorsal membrane ruffles upon tension drop

To examine the direct effect of membrane tension on PLD activity, we utilized hypo- or hypertonic buffer to induce the increase or decrease of membrane tension respectively in mouse myoblasts, similar to the approach taken by numerous prior studies (Boulant et al., 2011; Diz-Munoz et al., 2013; Houk et al., 2012). After transfection of a PA biosensor (PABD-GFP, <u>PA</u> <u>b</u>inding <u>d</u>omain of yeast Spo20p fused with GFP) into C2C12 myoblast, we imaged its distribution with inverted epi-fluorescence microscopy at 37 °C. Similar to a previous report (Zeniou-Meyer et al., 2007), PABD-GFP mainly distributed in nucleus and plasma membrane, especially at lamellipodia and membrane ruffles when cells were incubated in an isotonic buffer (1X PBS) (Fig. 1Aa). Interestingly, the enriched signals of PABD-GFP at plasma membrane became diffused after 2 min-incubation in a hypotonic buffer (0.5X PBS) (Fig. 1Ab), and it re-appeared when the cells were recovered by 1X PBS for 2 minutes (Fig. 1Ac). We hereafter referred this hypotonic treatment followed with isotonic buffer recovery as osmotic shock (OS) treatment.

A similar phenotype could be observed by directly incubating the cells with a hypertonic buffer (PBS + 150 mM sucrose) where PABD-GFP signal at membrane ruffles increased from 2 min, peaked at 5 min and gradually settled (Fig. 1B and S1A). We thus quantified the effect of hypertonic buffer treatment on membrane ruffling by dividing the ruffling phenotype into four categories: sparse, mild, intermediate and severe with <5%, 5-10%, 10-25% or >25% of dorsal area equipped with membrane ruffles respectively (Fig. 1C). While only 5% of cells incubated in isotonic buffer had vigorous membrane ruffles (intermediate and severe), 5-min hypertonic buffer incubation significantly increased this population to 52%. Together, we found that PABD-GFP signal at plasma membrane was enhanced when membrane tension was reduced by OS or hypertonic buffer treatment.

**Figure 1.**
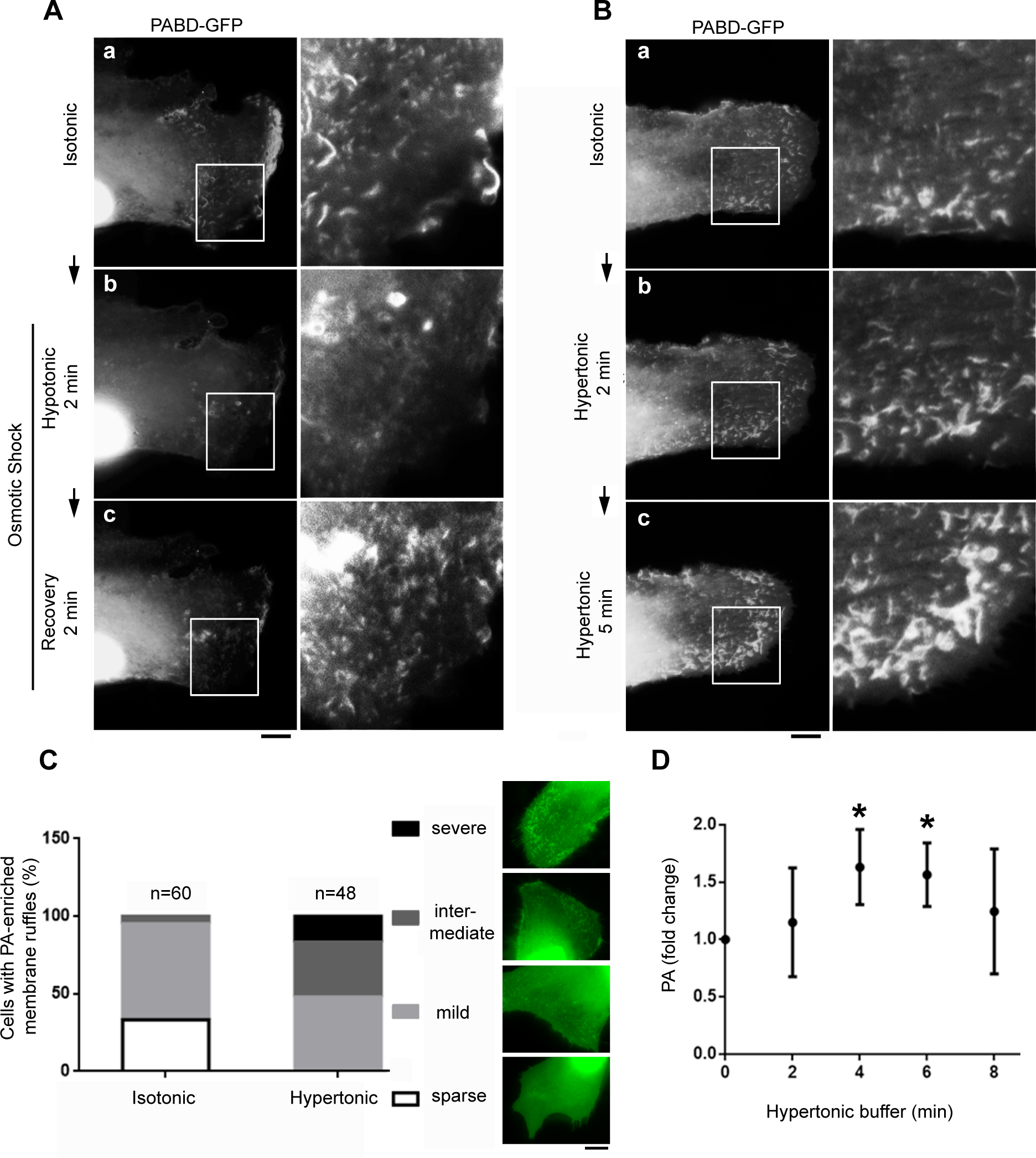
Effects of membrane tension manipulation on PA in myoblast. (A) An increase of membrane tension suppresses PA-rich ruffles at plasma membrane. PABD-GFP expressing C2C12 myoblast was first imaged in isotonic buffer (Aa, 1X PBS), and two images were further captured after 2 min of hypotonic buffer (Ab, 0.5XPBS) and additional 2 min of 1X PBS recovery (Ac). Boxed areas were magnified and shown on the right panel. (B) A decrease of tension induces PA enrichment at plasma membrane. PABD-GFP expressing myoblast was first imaged in isotonic buffer (Ba), and two images were further captured after 2 min (Bb) or 5 min (Bc) after hypertonic buffer treatment (1XPBS + 150 mM sucrose). (C) Quantification of membrane ruffling in cells treated with isotonic or 5-min hypertonic buffers. Ruffling phenotypes were divided into four categories with different levels or areas of membrane wave. (D) Kinetic change of total PA amount in myoblasts upon hypertonic buffer treatment. C2C12 myoblasts treated with hypertonic buffer for indicated time period were harvested and analyzed with total PA assay and normalized with the protein concentration. Fold change of the PA/protein ratio was compared with isotonic buffer treatment. Scale bars, 10 μm. *, *p* < 0.05.

To test whether the enrichment of PABD-GFP at plasma membrane reflects a change of cellular PA level upon tension alteration, we extracted total lipids from cells treated with different buffers, measured the amount of PA with an enzymatic assay, normalized with protein concentrations and expressed it as fold change compared with control cells. While the total PA amount in hypertonic buffer-treated cells showed similar kinetic accumulation with PABD-GFP signals which increased gradually and peaked at 5 min (Fig. 1D), all the tension manipulations (hypertonic, hypotonic and OS) led to increased total PA (Fig. S1B). These results suggest that total PA quantification may not reflect local PA production or distribution. To specifically investigate the effect of membrane tension on PA dynamics on plasma membrane, we thus mainly used PABD-GFP and live-cell imaging to further dissect this phenomenon.

To identify what the PA-rich membrane ruffles induced by tension drop are, we examined several plasma membrane components, including F-actin, dynamin-2 (Dyn2), phosphatydil-4,5-bisphosphate (PLC-PH-GFP, a probe for PI(4,5)P_2_), clathrin-coated pit adaptor protein (μ2-GFP) and PIP_3_ (labeled with Akt-PH-GFP) (Fig. 2A-F). In line with the function of PA on actin polymerization, we observed F-actin enrichment on the PABD-GFP ruffles where the membrane fission enzyme Dyn2 was also localized to (Fig. 2A, B). We thus used Dyn2-mCherry to mark the tension drop-induced membrane ruffles for other membrane components analysis between different pairs of markers. Interestingly, while PI(4,5)P_2_ was enriched and co-localized with Dyn2-mCherry, neither PIP_3_ nor μ2-GFP were enriched at those membrane ruffles (Fig. 2C-E). In addition to the enrichment of F-actin on membrane ruffles, dramatic accumulation of cortical actin and actin stress fibers also occurred upon hypertonic buffer treatment (Fig. 2F).

**Figure 2.**
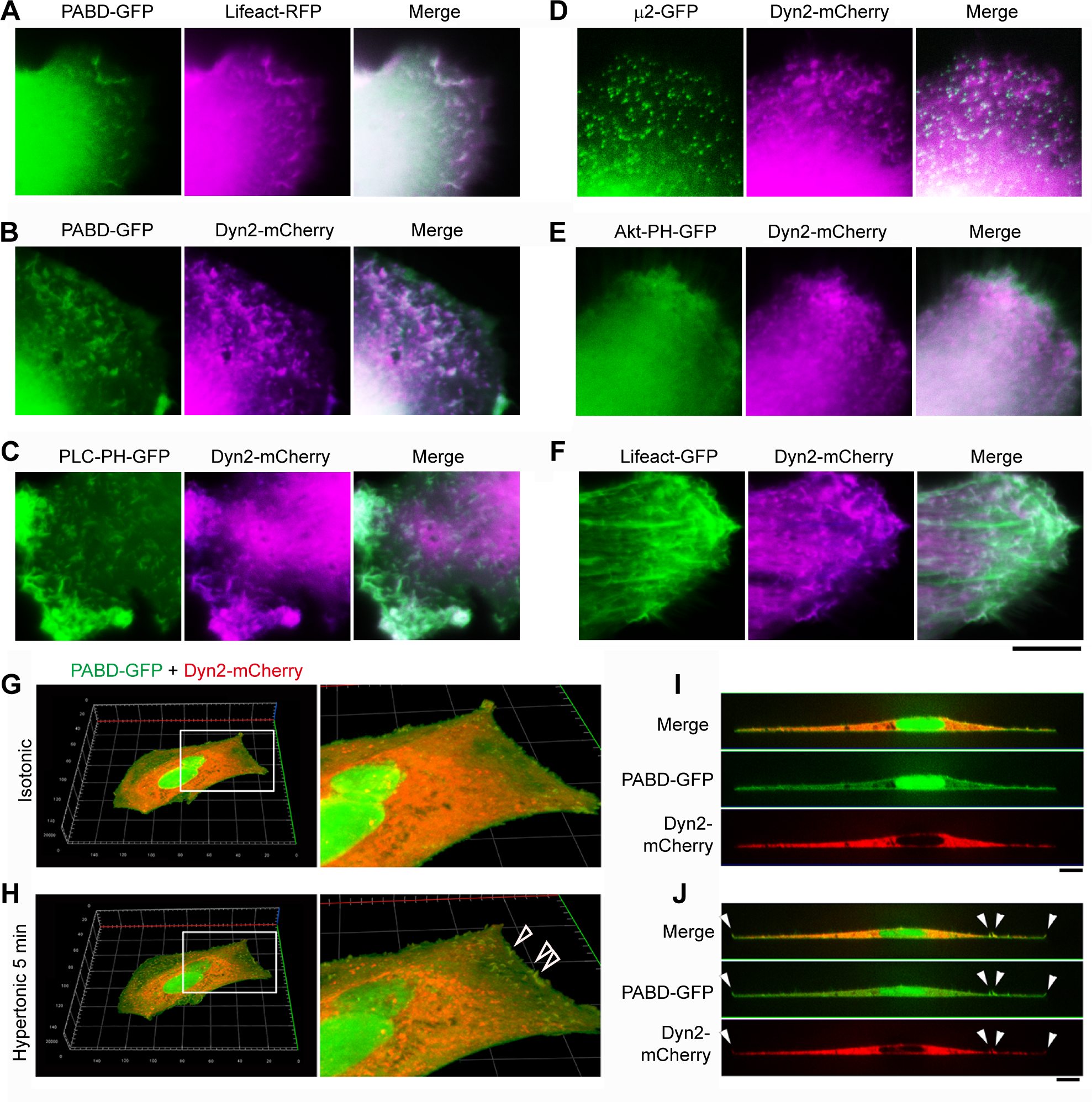
Distribution of membrane proteins and lipids upon sudden decrease of membrane tension. (A-F) Myoblasts co-transfected with indicated plasmids were treated with hypertonic buffer for 5 mins. The tension drop-induced PA-rich ruffles were examined for its enrichment with F-actin (Lifeact-RFP), Dyn2, clathrin-coated pit adaptor (µ2-GFP), PI(4,5)P_2_ (PLC-PH-GFP) and PI(3,4,5)P_3_ (Akt-PH-GFP). (G-J) Spinning disc confocal microscopy images of myoblast incubated in a hypertonic buffer. PABD-GFP and Dyn2-mCherry co-transfected myoblast was imaged with z-stack confocal microscopy before and after 5 min incubation in a hypertonic buffer. The 3D reconstructed images (G,H) and orthogonal views (I,J) were shown to illustrate dorsal membrane ruffling (open arrow heads in H and white arrow heads in J) upon tension drop. Scale bars, 10 μm.

To further visualize the membrane ruffles in three dimension (3D) when cell encounters acute tension reduction, we utilized spinning disk confocal microscopy to image the cell before and after hypertonic buffer treatment along the z-axis. After image acquisition and 3D reconstruction, we found that PABD-GFP and Dyn2-mCherry-enriched membrane ruffles were distributed at the peripheral dorsal membrane (open arrow heads in Fig. 2G, H). The decrease of cell volume upon hypertonic buffer incubation could be observed in the orthogonal image where there was a reduction in cell height without changing cell width (Fig. 2I, J). Notably, all the PA and Dyn2-enriched membrane ruffles existed only at the dorsal membrane. Together, we found that an acute decrease of plasma membrane tension results in F-actin and PA-enriched membrane ruffle formation at the peripheral dorsal membrane.

### An acute membrane tension drop induces macropinocytosis

The membrane flaccidity-induced F-actin and Dyn2-rich dorsal membrane waves were reminiscent of the membrane ruffles in macropinocytosis. To examine this possibility, we added a macropinocytosis-specific cargo, rhodamine-conjugated dextran (Rh-Dextran) into buffers with different osmotic conditions and incubated PABD-GFP expressing cells with these buffers. While no dextran signal was detected in cells incubated with isotonic buffer, several red puncta with about 1 μm in diameter were observed at the cell periphery in hypertonic and OS-treated cells together with prominent PABD-GFP ruffles (Fig. 3A). Quantification results showed that both OS and hypertonic buffer treatments result in ∼ 20% cells obtaining ≥ 3 dextran-positive macropinosomes (Fig. 3B). This result suggests that acute tension reduction induced by either direct hypertonic buffer incubation or OS treatment results in PA production, actin and Dyn2-rich membrane ruffling and subsequently macropinocytosis.

**Figure 3.**
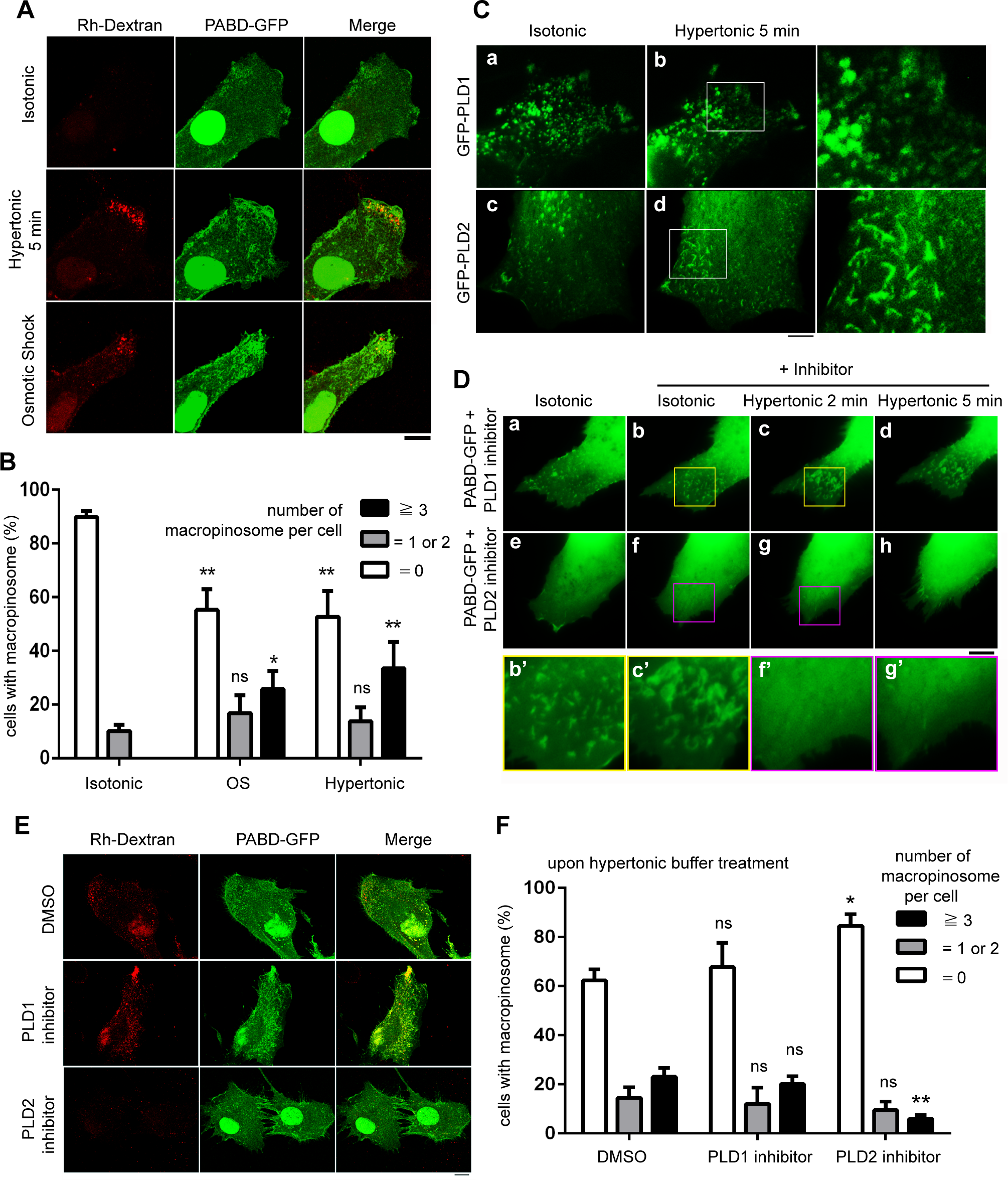
Acute membrane tension drop induces PLD2-dependent macropinocytosis. (A) Macropinocytosis is induced upon hypertonic buffer or OS treatment. PABD-GFP expressing myoblasts were incubated with indicated buffer plus 1 μg/ml Rh-Dextran, including 5 min isotonic (1X PBS), 5 min hypertonic (1XPBS + 150 mM sucrose) or 2 min hypotonic (0.5X PBS without dextran) followed with isotonic (1X PBS with dextran) for OS. After wash and fixation, cells were imaged with confocal microscopy and the maximum intensity projection images were generated. (B) The number of macropinosomes (Rh-Dextran containing vesicle larger than 0.5 μm in diameter) were quantified and divided into three populations as indicated. Percentage of each population were compared with the isotonic buffer-treated cells, and over 160 cells for each condition were analyzed. (C) PLD1 and PLD2 distribution upon membrane tension drop. Myoblasts expressing GFP-PLD1 or GFP-PLD2 were imaged before (Ca and Cc) and after (Cb and Cd) hypertonic buffer treatment. (D-F) Effect of PLD inhibitors on tension drop-induced membrane ruffles and macropinocytosis. PABD-GFP expressing myoblasts were pretreated with indicated PLD inhibitors, 500 nM for 10 min in 1X PBS, and followed by inhibitor-containing hypertonic buffer for another 5 min (D) or Rh-Dextran containing hypertonic buffer and OS treatment (E,F). Live cell imaging (D) or confocal imaging on fixed cells (E) were performed respectively. (F) The population of cells with different numbers of macropinosome were quantified with 150 cells of each condition and compared with vehicle control. Scale bars, 10 μm. *, *p* < 0.05; **, *p* < 0.01.

### PLD2 is responsible for PA production upon tension decrease

To examine whether PLD2 is responsible for the tension induced-PA production, we first used a general PLD inhibitor 1-butanol (Brown et al., 2007), to pre-treat myoblasts for 10 min and then incubated the cells with hypertonic buffer (Fig. S2). After 10 min incubation with 0.1% 1-butanol in isotonic buffer, mild cell contraction and diminished PABD-GFP signal on plasma membrane were observed (Fig. S2A). Further contraction of the cell was observed after hypertonic buffer treatment, yet no membrane ruffles were induced. By contrast, the control alcohol 2-butanol-treated cells showed prominent membrane ruffling enriched with PABD-GFP and Dyn2-mCherry upon tension drop. (Fig. S2B). This result indicates that PLD activity is essential for the PA-rich membrane ruffles induced by tension drop.

Mammalian cells express mainly two isoforms of PLD, PLD1 and PLD2, which are localized to intracellular membrane and plasma membrane, respectively. To examine the involvement of these two PLDs, we first monitored the distribution of PLD1 and PLD2 in cells treated with hypertonic buffer. While GFP-PLD1 remained intracellularly distributed in hypertonic buffer, GFP-PLD2 showed increased signals at dorsal membrane ruffles upon tension reduction (Fig. 3C). To further test the specific requirement of PLD2 activity upon tension drop, we used PLD1 or PLD2 specific inhibitors, UV0359595 and UV-364739 respectively, to pre-treat the cells for 10 min in 1X PBS and followed by 5-min hypertonic buffer incubation in the presence of inhibitor. Consistent with the subcellular localization results, we found that only PLD2 inhibitor blocked the membrane ruffling induced by tension drop, but not in PLD1 inhibitor-treated cells (Fig. 3D). Consistent with the data above, only the PLD2 inhibitor blocked Rh-Dextran internalization induced by hypertonic buffer incubation (Fig. 3E, F).

To substantiate the specific requirement of PLD2 in tension drop-induced macropinocytosis, we knocked down (KD) either PLD1 or PLD2 with two different lentiviral shRNA sequences. Similar with the results of PLD inhibitors, we observed significant reduction of membrane ruffles and macropinosome formation in PLD2 KD cells, not PLD1 KD cells (Fig. 4A-C). It is worth noting that when PLD1 or PLD2 were knocked down, the expression of the other isoform was slightly increased (Fig. 4D), indicating a compensatory expression and adaptation phenomenon of cells with PLD depletion.

**Figure 4.**
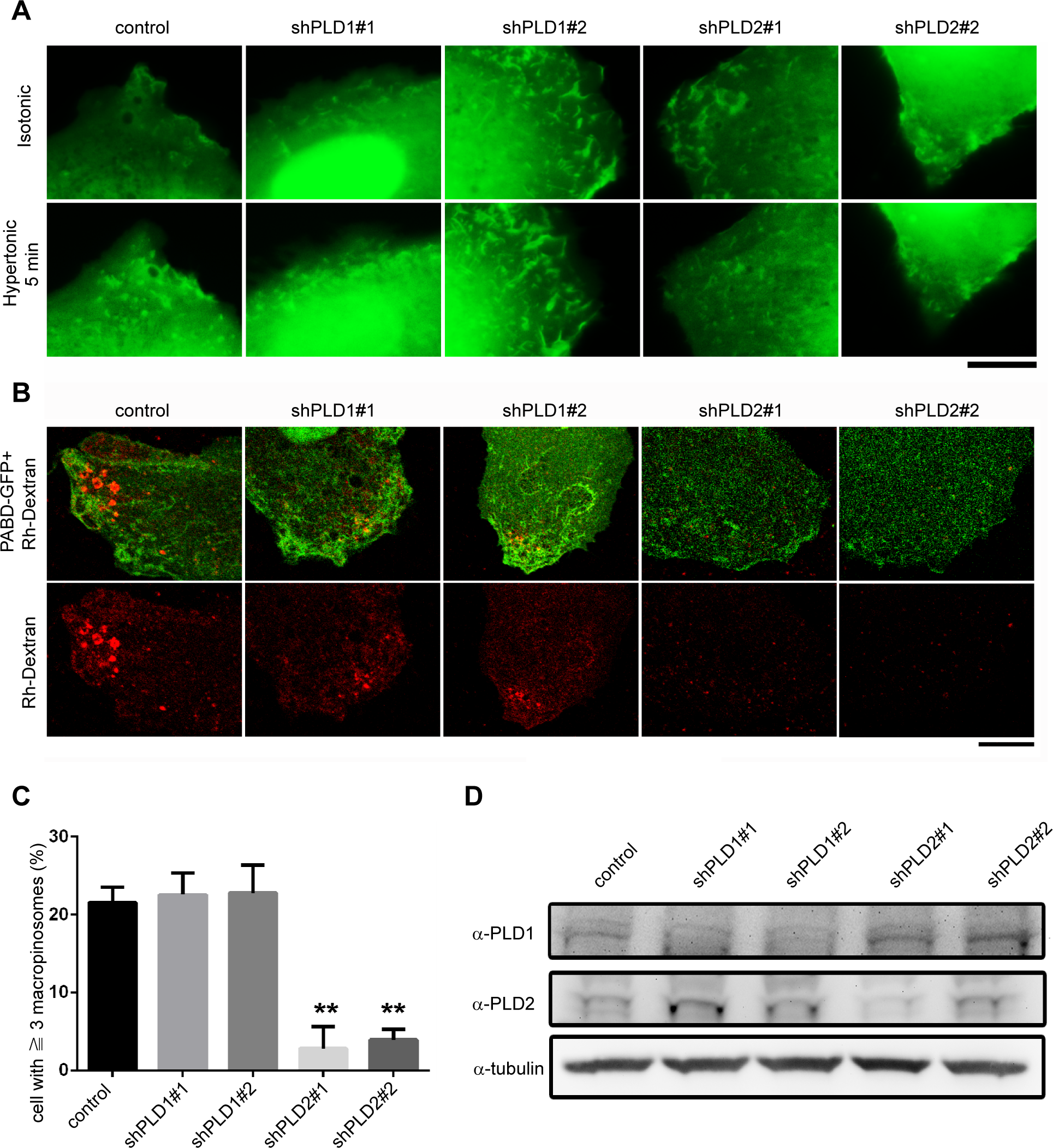
PLD2 is required for tension drop-induced membrane ruffling and macropinocytosis. (A) PABD-GFP transfected control, PLD1- or PLD2-depleted myoblasts were subjected to hypertonic buffer treatment, and the distribution of PABD-GFP on dorsal membrane was captured by inverted fluorescence microscopy. (B) Macropinocytosis of PABD-GFP myoblasts upon hypertonic buffer treatment were monitored with Rh-Dextran internalization in cells PLD1 and PLD2 depleted cells compared to control. (C) The population of cells with more than three macropinosomes (the diameter ≥ 0.5 μm) as determined by Rh-Dextran for experimental conditions in (B) were quantified and compared with control cells. 120 cells of each condition were scored. (D) Western blotting of PLD1 and PLD2 in myoblast upon PLD shRNA KD. Scale bars, 10 µm. **, *p* < 0.01.

Together, these results suggest that membrane flaccidity induces PA production and macropinocytosis *via* PLD2 activity. Notably, the intracellular GFP-PLD1 vesicles were diminished upon tension surge concurrent with the increased GFP-PLD1 signals on plasma membrane (Fig. S3A), whereas GFP-PLD2 was less enriched on plasma membrane with hypotonic buffer treatment (Fig. S3B). These results indicate that both PLDs respond to tension change and may explain why both an increase and a decrease of membrane tension caused the increase of total PA amount (Fig. S1B).

### Lipid microdomain responds to tension alteration

Lipid raft has been reported as a mechanosensor and could be disrupted chemically or mechanically (Oglecka et al., 2014; Petersen et al., 2016). We wondered whether the integrity of lipid microdomain may be affected by membrane tension *in vitro* and *in vivo*. To directly observe the effect of membrane tension on lipid microdomain, we utilized giant unilamellar vesicles (GUVs), a commonly used membrane template for studying lipid phase separation, to monitor the size of different lipid phases with L_o_ (liquid-ordered) domain labeled with Topfluor cholesterol and L_d_ (liquid-disordered) domain labeled with rhodamine-PE. In GUVs composed of DOPC:DPPC:Cholesterol:Topfluor cholesterol: rhodamine-PE with molar ratio at 39:39:20:1:1, we observed phase separation with several L_o_ domains (around 3 – 15 per GUV) when the GUVs were incubated in isotonic buffer (Fig. 5A and video 1). Strikingly, when GUVs were incubated in hypotonic buffer, those L_o_ domains merged into one large domain with numerous small L_o_ domains continuously formed and fused together (Fig. 5A and video 2). By contrast, L_o_ domains became smaller in hypertonic buffer where GUVs on a coverslip resemble flatten tires (Fig. 5A and video 3). These results convincingly showed that membrane tension affects the integrity of lipid microdomain on model membrane.

**Figure 5.**
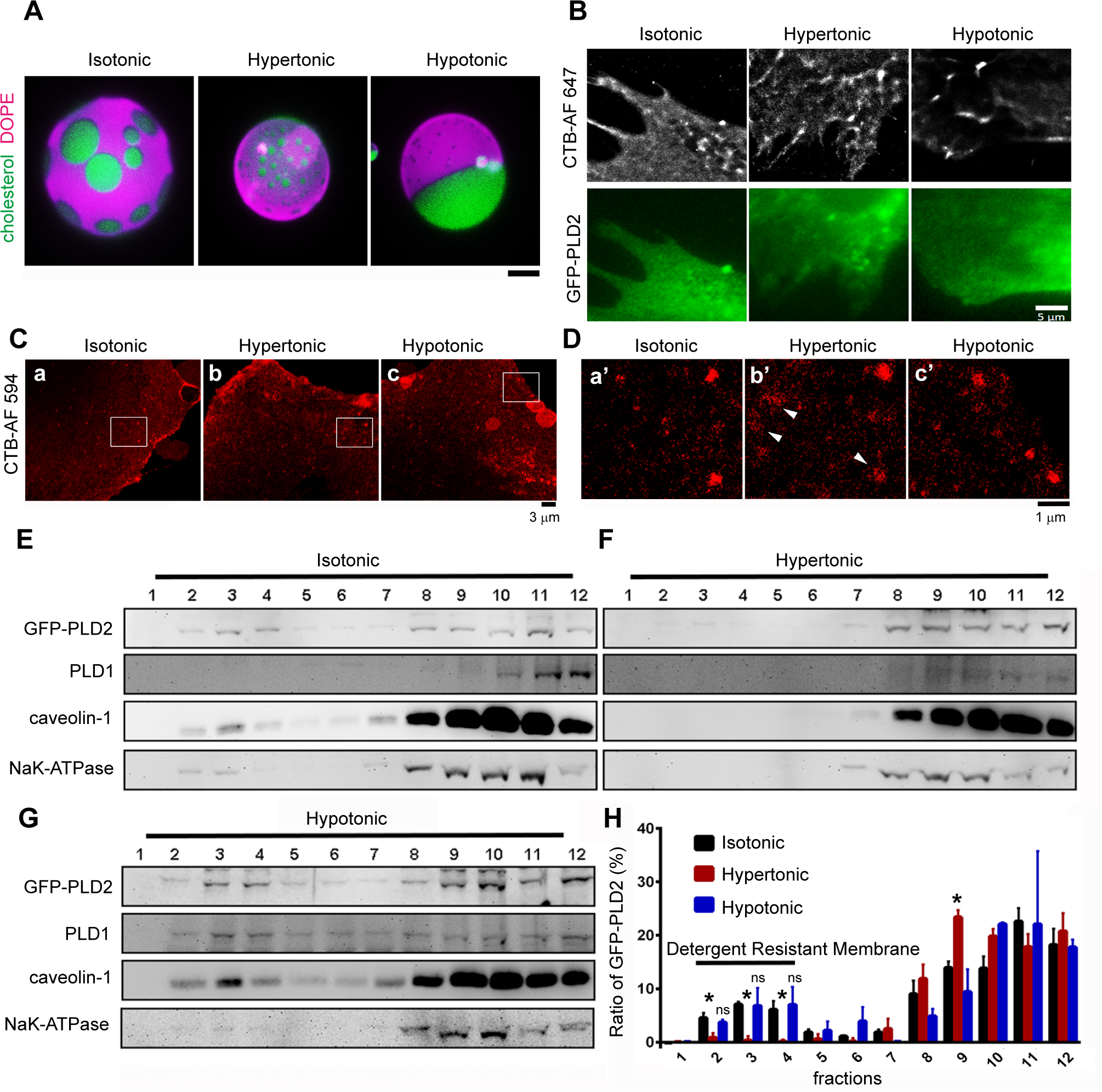
Lipid microdomain disruption upon tension drop. (A) Effects of tension changes on lipid phase separation. GUVs with Topflour cholesterol (green) and rhodamine-PE (magenta) to label L_o_ and L_d_ domains respectively were subjected to iso- (400 mM), hyper- (500 mM), or hypo-tonic (300 mM) glucose buffers and imaged with inverted epi-fluorescence microscopy. Scale bar, 10 μm. (B,C) Effects of hyper- or hypotonic buffer treatment on lipid rafts in cell. Myoblasts were treated with indicated buffers, fixed, stained with CTB and imaged with epi-fluorescence (GFP-PLD2) or STORM (CTB-AF 647) (B) or STED microscopy (C), respectively. (D) Boxed areas in (C) were magnified. (E-G) Fractionations of detergent-resistant membrane proteins upon tension changes. GFP-PLD2 expressing myoblasts treated with indicated osmotic buffers were harvested and lysed with buffer containing 1% Triton X-100 at 4 °C for 30 min and subjected to sucrose gradient ultracentrifugation. Detergent resistant membranes were distributed in fraction 2-4. (H) The ratio of GFP-PLD2 in each fraction was analyzed with Student *t*-test. Three repeats of the experiments were quantified and compared with isotonic buffer-treated cells. *, *p* < 0.05.

To test whether the flaccidity-induced disruption of lipid microdomain could be observed in a cell, we utilized Alexa flour 647-conjugated cholera toxin B (CTB) which binds to sphingolipid GM1 to label lipid raft and monitored its distribution together with GFP-PLD2. Upon different osmotic buffer treatments, fixation and CTB-AF 647 staining, images of dorsal membranes were acquired with Stochastic Optical Reconstruction Microscopy (STORM) and analyzed with ThunderSTORM plugin in ImageJ (Fig. 5B). We observed partial co-localization between GFP-PLD2 and CTB labeled lipid microdomains in hypertonic buffer-treated cells; however, the size of CTB-labeled microdomains were similar among the three conditions. We further used Stimulated Emission Depletion (STED) Microscopy to image the ventral surface of myoblasts after indicated osmotic buffer treatments and found no significance difference in the size of CTB-AF 594 puncta in different buffer-treated cells (Fig. 5Ca-c), though the CTB clusters seemed less compact in hypertonic buffer-treated cells (arrow heads in Fig. 5D).

To quantitatively examine the effect of tension change in cells, we utilized nonionic detergent to isolate detergent-resistant membrane under different tension conditions (Lingwood and Simons, 2007). Given the weak signals of endogenous PLD2 in C2C12 myoblast (Fig. 4D), we ectopically expressed GFP-PLD2 in myoblast, treated the cells with buffers with distinct osmolarity and isolated the detergent-resistant membrane with 1% cold TX-100 and sucrose gradient ultracentrifugation, then detected the distribution of interested proteins with Western blotting (Fig. 5E-H). In addition to GFP-PLD2, we also detected other membrane proteins, including PLD1, caveolin-1 and Na^+^-K^+^ ATPase for comparison. Both GFP-PLD2 and caveolin-1 became less resistant to detergent (less enrichment in fractions 2-4) in cells at a lower tension (Fig. 5F), supporting the idea that tension drop destabilized lipid rafts. In contrast, lipid rafts remained intact in cells with increased membrane tension (Fig. 5G,H). Interestingly, PLD1 partially shifted to detergent-resistant membrane fractions in hypotonic condition that is consistent with its partial redistribution to plasma membrane upon tension surge (Fig. 5G and S3A). Together, these results demonstrate that membrane tension facilitates lipid phase separation, thus a decrease of tension results in lipid raft destabilization *in vitro* and *in vivo*.

### Acute decrease in membrane tension in myotube induces macropinosome formation

Given the results above, we have identified a previously underappreciated pathway which transduces mechanical cue into biochemical signal, *i.e.* the membrane tension alteration leads to phosphatidic acid production. However, compared to the pronounced actin polymerization (Fig. 2F), we reasoned the relatively small amount of membrane ruffles and macropinosome in C2C12 myoblast upon tension drop indicates that PA-induced macropinocytosis contributes to a limited extent for cell to respond to tension alteration. We thus wonder whether this pathway will be more pronounced in cells with restricted actin dynamics, for example, in the muscle cells.

Skeletal muscle is constantly exposed to mechanical stress in our body thus has been intensively studied for its ability to cope with increased membrane tension. Muscle cell is equipped with robust caveolae structures which could quickly disassemble when membrane tension increases in order to release membrane reservoir thus relieve the tension surge (Lo et al., 2015; Sinha et al., 2011). However, little is known about how muscle cells react upon rapid membrane tension drop following a tension surge. After hypertonic buffer or OS treatment in the presence of Rh-Dextran, we observed significant amounts of dextran-containing macropinosomes in C2C12-derived myotubes (Fig. 6A). Remarkably, the macropinosomes induced by OS were much larger than the ones in hypertonic buffer-treated myotubes which could be observed under light microscopy (Fig. 6Bc, red arrow heads). This is probably due to the increase of membrane area by caveolae disassembly during hypotonic treatment (black arrow in Fig. 6Bb) thus results in more pronounced macropinosomes than cells in hypertonic buffer incubation. Furthermore, using time-lapse microscopy, we found those OS-induced, dextran-positive macropinosomes were also enriched with PA (Fig. 6C).

**Figure 6.**
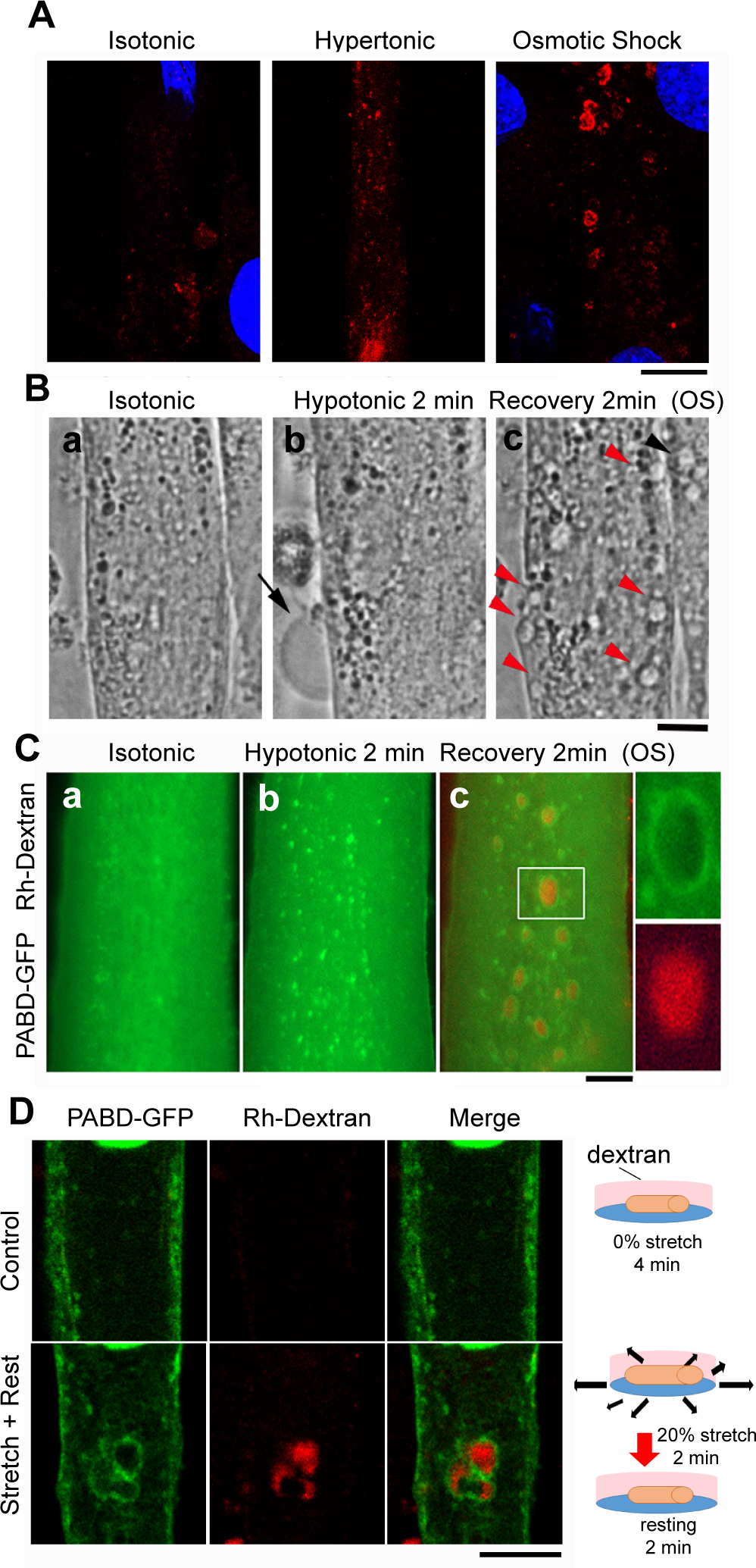
Acute tension drop induces macropinocytosis in myotubes. (A) Macropinosome formation in myotubes. C2C12-derived myotubes incubated in indicated buffers plus Rh-Dextran. After intensive wash, cells were fixed, stained with DAPI and imaged with confocal microscopy. (B) Morphology of myotubes treated with isotonic buffer (Ba), hypotonic buffer (Bb) and followed by isotonic buffer recovery (Bc). Phase contrast microscopy images were captured with inverted fluorescence microscopy. Black arrow indicates the membrane bleb from cell swelling, and the red arrow heads point to macropinosomes induced by tension drop. (C) Macropinosomes are enriched with PA in OS-treated myotube. Live cell imaging of PABD-GFP expressing myotube was captured when cells were incubated in isotonic buffer (Ca), hypotonic buffer (Cb) and recovery isotonic buffer containing 1 μg/ml Rh-Dextran for 2 min with intensive wash (Cc). Insets are magnified images from the boxed areas. (D) Radial stretch induces macropinosome formation in myotubes. PABD-GFP expressing myotubes incubated in 1 μg/ml Rh-Dextran medium were subjected to 0% or 20% radial stretch for 2 min and followed by 2 min resting (0% stretch). After intensive wash with PBS, myotubes were fixed and imaged with confocal microscopy. Scale bars, 10 µm.

To examine whether macropinocytosis induced by OS treatment is physiologically relevant, we stretched and relaxed PABD-GFP expressing myotubes with radial cell stretcher for 2 min-stretching and 2 min-relaxation in the presence of Rh-Dextran in medium. After wash and fixation, cells were imaged with confocal microscopy and we observed some macropinosomes labeled with dextran and PABD-GFP in myotubes, but not in the control myotubes which were incubated in dextran without stretching (Fig. 6D). These results demonstrate that acute membrane tension reduction induced by intensive cell stretching and relaxation, similar to OS treatment, triggers PA production and macropinocytosis. We hereafter utilized OS-treated myotubes to observe the kinetic and PLD contribution to tension change-induced macropinocytosis in myotube.

In contrast to what we observed in myoblast, there was no PA-rich membrane ruffles in myotubes upon OS treatment (Fig. 6B,C). Thus, we carefully examined the kinetic distribution of Dyn2-mCherry and PABD-GFP in myotube treated with OS (Fig. 7A). Although there was no membrane ruffling detected in OS-treated myotubes, we found Dyn2-mCherry enriched at PABD-GFP ring at membrane during the early stage of OS treatment (Fig. 7A, inset). Interestingly, only when the macropinosome was formed above the nucleus where much less cytosolic signals exist that the Dyn2-mCherry could be observed on the membrane, potentially the intermediate stage of macropinocytosis (Fig. 7A). In addition, different from the PA-labeled macropinosomes that could last for more than 10 min in myotubes, the Dyn2-mCherry ring quickly disappeared from plasma membrane upon OS treatment, suggesting the disappearance of Dyn2 or the disassembly of Dyn2 ring after membrane scission (Antonny et al., 2016). Together, these results suggest that an acute membrane tension decrease also induced PA and Dyn2-mediated macropinocytosis in myotube though without displaying a membrane ruffling phenotype.

**Figure 7.**
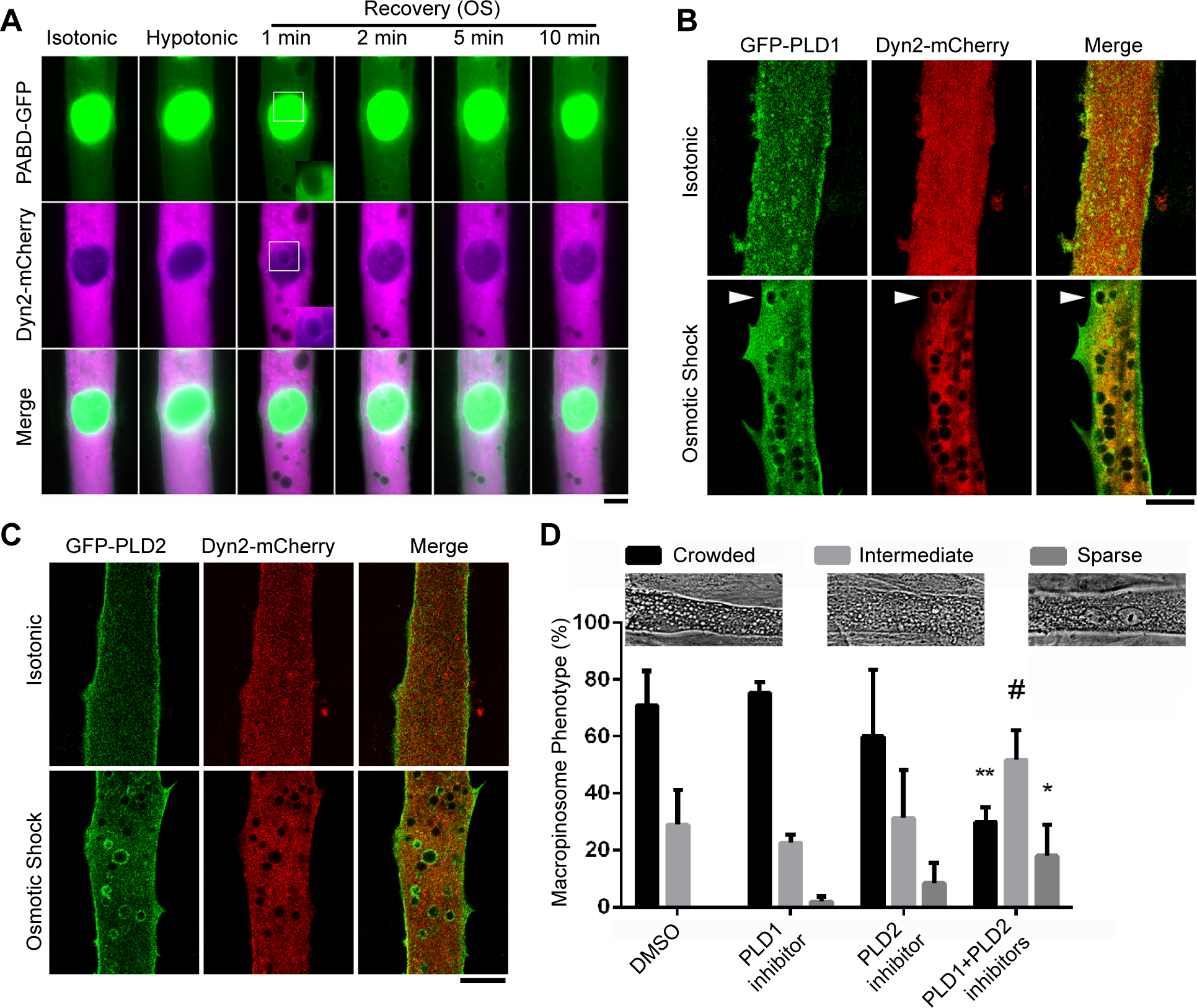
Both PLD1 and PLD2 contribute to the macropinosome formation in OS-treated myotubes. (A) Kinetic distribution of PABD-GFP and Dyn2-mCherry in an OS-treated myotube. Boxed areas were magnified and adjusted for lower green fluorescence intensity to better observe the macropinosome on top of a nucleus. (B, C) Distribution of PLD1 and PLD2 in OS-induced macropinosomes. C2C12-derived myotubes expressing GFP-PLD1, GFP-PLD2 and Dyn2-mCherry were treated with isotonic buffer or OS and followed by fixation and image acquisition with confocal microscopy. Single focal images were shown with scale bars of 10 μm. (D) Effects of PLD inhibitors on OS-induced macropinosome formation. Myotubes were pre-treated with indicated inhibitors for 10 min followed by OS that also contained the indicated inhibitors. Phase contrast images of treated cells were acquired with inverted microscopy and the phenotypes were divided into three categories: crowded (macropinosome with diameter > 2 μm), intermediate and sparse as shown in graph. Data from three independent experiments with n > 100 myotubes for each condition were compared to DMSO-treated control. #, *p* < 0.1; *, *p* < 0.05; **, *p* < 0.01.

Next, we asked whether PLD is also responsible for the tension change-induced macropinocytosis in myotube. Interestingly, PLD1-GFP distributed both on plasma membrane and intracellular vesicles in myotube and partially distributed to macropinosomes after OS, whereas PLD2 was localized at the plasma membrane in isotonic condition and became enriched at macropinosomes upon OS treatment (Fig. 7B,C). Consistent with these observations, OS-induced macropinosome formation in myotubes could only be blocked by the combined treatment of PLD1 and PLD2 inhibitors (Fig. 7D). Together, these data indicate that the tension drop-PLD-macropinocytosis pathway is dominant in myotubes and both PLD1 and PLD2 are involved.

### PA enhances the membrane fission activity of Dyn2

PA is known to promote actin polymerization through PI(4,5)P_2_ production and Rac activation (Liu et al., 2013). However, it has not been addressed whether PA could directly affect the activity of Dyn2 which is a membrane fission GTPase essential for phagocytosis and macropinocytosis. (Liu et al., 2008; Marie-Anais et al., 2016). Furthermore, the fission activity of Dyn2 is tightly regulated by membrane curvature and lipid composition (Morlot et al., 2012; Pucadyil and Schmid, 2008; Roux et al., 2010). To test whether PA could facilitate the activity of Dyn2, we first performed liposome sedimentation assays to examine the membrane binding ability of purified human Dyn2 to liposomes containing different amount of PA. There was no significant difference of Dyn2 binding to PA-containing liposomes (Fig. S4A, B). Interestingly, the GTPase activity of Dyn2 increased in a PA-dependent manner (Fig. S4C). We further analyzed the assembly of Dyn2 on liposomes and its membrane fission activity with negative stain electron microscopy and *in vitro* membrane fission assay. We found that not only did Dyn2 assemble into much ordered and distinct helical structures in PA-containing liposomes (Fig. S4D), it also showed higher fission activity on PA-containing membrane template (Fig. S4E). Interestingly, the fission activity of Dyn2 seemed lower on 10% PA membrane compared to 5% PA membrane which is probably due to the instability and spontaneous membrane breakage on membrane reservoir made with high amount of PA. Together, PA facilitates the proper assembly of Dyn2 on membrane without changing the fraction bound, thus enhances its GTP hydrolysis and membrane fission activities.

## Discussion

Macropinocytosis is a unique endocytic pathway which is initiated by biochemical stimulations to efficiently uptake nutrients, terminate signaling or internalize viruses (Buckley and King, 2017). In this study, we identified a PLD-dependent, but PIP_3_-independent pathway of macropinocytosis that is induced by acute membrane tension drop which leads to lipid raft destabilization, PLD2 activity, PA production and macropinosome formation (Fig. 8). This particular mechanotransduction pathway contributes differentially among cell types and is much pronounced in myotubes and reflects the molecular mechanism of muscle cell coping with intensive stretching and relaxation during physical exercise.

**Figure 8.**
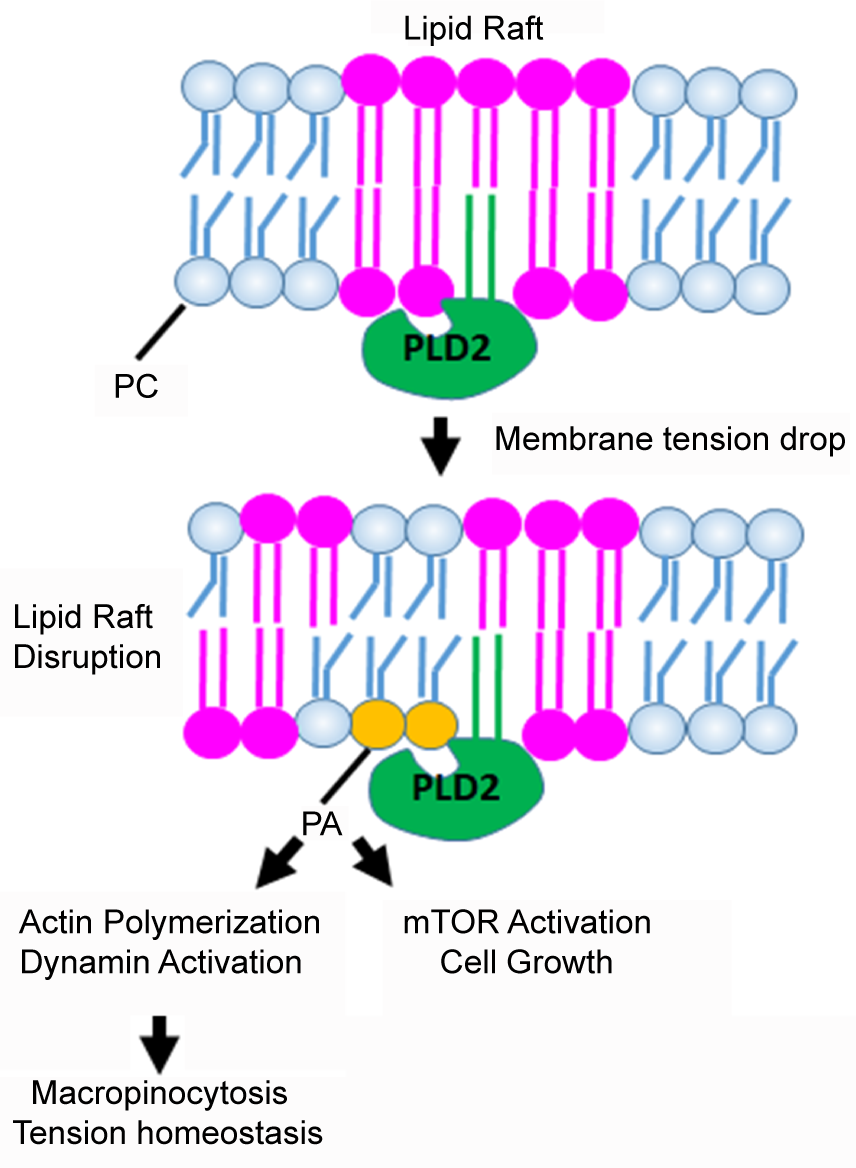
Model of tension drop-induced lipid raft disruption and macropinocytosis. Under normal tension, lipid phase separation results in lipid-ordered domain formation thus segregates PLD2 from its activator PI(4,5)P_2_ as well as its substrate PC (upper panel). When membrane tension is reduced suddenly, the decreased line tension would favor smaller lipid microdomains thus allow PLD2 to access its activator and substrate. The production of PA on plasma membrane leads to PI4P-5 kinase activation to generate PI(4,5)P_2_ and promotes higher PLD activity, actin polymerization, Dyn2 activation and subsequently membrane ruffling and macropinocytosis.

Lipid raft is well perceived as a signaling platform given its enrichment of unique receptors. Here we find it also serves as a mechanosensor which undergoes disruption upon membrane tension alteration to activate PLD2. The effect of membrane tension of the coarsening of lipid microdomains *in vitro* has been largely studied and modeled theoretically (Akimov et al., 2007; Chen and Santore, 2014; Hamada et al., 2011; Oglecka et al., 2014; Portet et al., 2012; Ursell et al., 2009). It is believed that the membrane tension sensitivity of lipid microdomain may arise from (1) the increase of line tension when membrane tension rises (Akimov et al., 2007), or (2) the kinetics of microdomain coalescence affected by membrane tension (Ursell et al., 2009). Line tension is the boundary energy between L_o_ and L_d_ domains that comes from the difference of their membrane thickness, also known as hydrophobic mismatch. Therefore, it is intuitive to imagine that membrane with lower tension would have lower boundary energy thus the total circumference of L_o_ could increase, *i.e.* a larger number of L_o_ domains with smaller diameters (Fig. 8). With that, it is tempting to hypothesize that PLD2 catalyzes reactions at the boundary of lipid rafts and the effect of membrane tension on lipid raft is to increase the amount of phase boundaries which leads to higher PLD2 activity.

The activity of PLDs has been reported to be sensitive to mechanical stress such as mechanical stretching, membrane tension surge or osmotic stress (Diz-Munoz et al., 2016; Hornberger et al., 2006; Tomassen et al., 2004). However, how exactly PLD is activated upon mechanical stimulation remains unclear. Here, we found that PLD1 and PLD2 distribute differently and react to distinct tension alterations in myoblasts. Nonetheless, it is puzzling that both an increase and a decrease of membrane tension result in elevated total amount of PA, yet resulted in different cellular outcomes: actin depolymerization or polymerization, respectively (Diz-Munoz et al., 2016; Tsujita et al., 2015). Since the subcellular location of PA and its surrounding proteins determine where and what kind of cellular event it would participate (Colley et al., 1997; Du et al., 2004; Teng et al., 2015; Yang and Frohman, 2012), it is plausible that elevated PA in different membrane compartment would trigger distinct cellular processes. Furthermore, given that high membrane tension directly inhibits actin polymerization and the force feedback mechanism on actin assembly (Bieling et al., 2016; Houk et al., 2012; Tsujita et al., 2015), the effects of membrane tension on PA production are important contributors to the final phenotype outputs. Further studies are needed to elucidate the feedback regulation of membrane tension, local PA production and actin polymerization in order to fully deciphering the complexity of this network.

PA has a small phosphate head group thus could create lipid packing defect in membrane by its nature (Bigay and Antonny, 2012). Here we found that membrane with PA facilitates the assembly of Dyn2, but not its binding, which is reminiscent of the effect of membrane curvature on Dyn2 assembly and activity (Liu et al., 2011a; Roux et al., 2010). Curved membrane provides more lipid packing defect thus facilitates the insertion of the PH domain in Dyn2 to have better assembly. With that, membrane with more PA may have more lipid packing defects due to its geometry. Consistent with this notion, PA has been reported to induce the insertion of Dyn1 into lipid monolayer (Burger et al., 2000). Together, these results suggest that Dyn2 assembly is better positioned for its fission activity on PA-containing membrane without the prerequisite of membrane curvature.

The mechanotransduction pathway we discovered here supports the concept that membrane tension is an integrator for biochemical and physical cues to regulate membrane trafficking and actin organization. By examining the molecular basis of this regulatory network, we uncovered the mechanisms underpinning membrane organization and PLD activity. Given the ubiquitous nature of forces, we suspect there are more molecules that sense and/or are regulated by membrane tension. Our findings pave the way for future studies of the mechanical regulation on membrane and cell physiology.

## Materials and Methods

### Cell Culture, Transfection and Infection

Mouse-derived C2C12 myoblasts (American Type Culture Collection, CRL-1772) were cultured in growth medium (GM), DMEM supplemented with 2 mM L-glutamine, 1 mM sodium pyruvate, antibiotics and 10% fetal bovine serum (Gibco). To induce differentiation, C2C12 were seeded onto laminin (Invitrogen)-coated glass-bottom dish (MatTek) or coverslips in GM, grown to 90% confluency, and then switched to differentiation medium (DM), which is the same as GM but with 2% horse serum (Gibco). This time point was considered as day 0 of differentiation. For transfection, cells at 70% confluency were transfected with the desired DNA constructs using Lipofectamine 2000 (Invitrogen), as recommended by the manufacturer.

For PLD knockdown experiments, lentiviruses with shRNA sequences: 5’-CCCAATGATGAAGTACACAAT-3’ or 5’-GCTTGGTAATAAGTGGATAAA-3’ for PLD1; whereas 5’-CCTTCCTGTCACCAAGTTCAA-3’ or 5’-CATGTCTTTCTATCGCAATTA-3’ for PLD2, were prepared. C2C12 myoblast were infected with control or PLD-shRNA containing lentiviruses and selected with 2 μg/ml puromycin for 3 days then subjected to transfection and osmotic buffer treatments.

### Reagents

PLD1 inhibitor UV0359595 was purchased from Cayman Chemical. PLD2 inhibitor UV0364739 was from Tocris Bioscience. Rh-dextran, CTB-AF 647 and CTB-AF 594 were from Invitrogen. Anti-PLD2 antibody was purchased from Abcam, anti-PLD1 and anti-caveolin-1 antibodies were from Cell Signaling Technology, anti-Na,K-ATPase antibody was from Santa Cruz Biotechnology, and anti-GFP antibody was a gift from Fang-Jen S. Lee in National Taiwan University. All lipids were purchased from Avanti Polar Lipids.

### Microscopy

For live-cell microscopy, cells transfected with interested DNA constructs were seeded on glass-bottom dish (MatTek) and imaged with Zeiss inverted microscopy Axio Observer Z1 or spinning disc confocal microscope (Carl Zeiss Observer SD) at 37 °C with 63×, 1.35-NA oil-immersion objective. To image fixed cells, sample slides were observed with confocal microscopy LSM700 with 63×, 1.35-NA oil-immersion objective (Carl Zeiss, Jena, Germany).

For super-resolution microscopy, we used both STORM and STED to image cells upon hyper- or hypo-osmotic shock. For STORM, GFP-PLD2 transfected C2C12 myoblasts were subjected to iso- (1X PBS), hypo- (0.5X PBS), or hypertonic (1X PBS with 150 mM sucrose) buffers for 5 minutes. After 4% paraformaldehyde fixation and intensive PBS wash, cells were incubated with Alexa fluor 647-cholera toxin B (CTB, 1 µg/ml, Invitrogen) overnight at 4 °C. After PBS wash, the STORM buffer (50 µL beta-mercaptoethanol (BME), 1.85 mM monoethanolamine (MEA), 0.25 M NaCl, 50 mM Tris with a final pH of 9.0) was added with protocatechuic acid (PCA)/protocatechuate-3,4-dioxygenase (PCD) O_2_-scavenging system immediately before imaging was used for switching the fluorophores. The imaging was performed at the Single Molecule Analysis in Real-Time (SMART) Center, University of Michigan, using computer-controlled IOX-81 Olympus microscope with 60× oil 1.49NA (APON60XOTIRFM) and an Andor iXon Ultra EM-CCD camera. 10000 images of Af 647-CTB excited with 640 nm (Coherent Cube) laser were collected at 100 ms exposure time using stream acquisition. After acquisition of 5000 images with 640 nm laser illumination, sample was co-exposed to 405 nm (Coherent Cube) laser to excite fluorophores to triplet state. Samples were exposed to both 640 nm and 405 nm until acquisition of 10000 images. A GFP-PLD2 image was taken before and after the STORM imaging. The STORM image reconstruction was performed using ThunderSTORM plugin in ImageJ, using difference of Gaussian image filtering and sub-pixel localization of molecules using maximum likelihood fitting with integrated Gaussian PSF.

For STED imaging, cells treated with different osmotic buffers and followed by fixation and Alexa fluor 594-CTB (Invitrogen) staining were mounted in ProLong Gold antifade reagent (Invitrogen) and imaged under Leica TSC SP8 X STED 3X with 100× oil objective 1.4NA (STED). Samples were acquired with excitation laser 594 nm, 660 nm depletion laser and Hybrid Detector (Leica HyD).

### PA measurement

Total PA content was measured with a coupled enzymatic reaction assay (Total Phosphatidic Acid Assay Kit, Cell Biolabs, Inc.). Briefly, cells treated with buffers of different osmolarities were scraped from dishes. While 10% of the samples were used for protein concentration analysis, the rest of the samples were subjected to total lipid extraction by methanol and chloroform. After drying with SpeedVac, the lipid film was dissolved and the amount of PA was determined by a fluorometric assay that firstly hydrolyzed PA to glycerol-3-phosphate by lipase. Next, glycerol-3-phosphate product was oxidized by glycerol-3-phosphate oxidase, producing hydrogen peroxide which reacted with a fluorometric probe with excitation/emission wavelength at 530-560 nm/585-595nm. The PA amount was determined as fluorescence intensity (A.U.)/protein concentration (µg).

### Macropinocytosis assay

To monitor macropinocytosis, myoblasts or myotubes were incubated with indicated osmotic buffers containing 1 µg/ml Tetramethylrhodamine Dextran (Thermo Fisher Scientific). After PBS wash for five times, cells were fixed and imaged with confocal microscopy. To monitor macropinocytosis in myotube upon cell stretching, myoblasts were seeded on laminin-coated flexcell silicon dish and were induced to differentiate for five days after PABD-GFP transfection. Day-5 differentiated myotubes were incubated in DM medium containing 1 µg/ml Rh-Dextran and subjected to radial stretching of 2 min at 20% extension and followed by 2 min resting. After intensive wash, fixation and mounting, cells were imaged by confocal microscopy.

### Fractionations of detergent-resistant membrane domains

Detergent-resistant membrane domains were purified from cells exposed to buffers with different osmolarities as previously described (Czarny et al., 1999). Briefly, after incubation with buffers of different osmolarity, myoblasts (two sub-confluent 100-mm dishes) were scraped into ice-cold buffers with the same osmolarity as previous treatment. After centrifugation, the pelleted cells were re-suspended with 0.8 ml lysis buffer (50 mM Tris, pH 7.4, 150 mM NaCl, 2 mM EDTA, 1% Triton X-100 and proteinase inhibitor cocktail (Roche)). After 30 min incubation at 4 °C, cell extracts were adjusted to 40% sucrose by addition of 0.8 ml of the above buffer (minus Triton X-100) containing 80% sucrose and is then placed at the bottom of a 5-ml ultracentrifuge tube. A step sucrose gradient was formed above the lysate by adding 2.4 ml of 35% and 0.8 ml of 5% sucrose solutions, and the tubes were centrifuged at 150,000 x*g* for 8 h in a SW-55 Ti rotor at 4 °C. Fractions (0.4 ml) were collected from the top of the gradient and analyzed with Western blotting.

### Liposome and GUV preparation

For liposomes preparation, lipid mixtures (DOPC:DOPS:PI(4,5)P_2_:DOPA at 80-70:15:5:0-10) were dried, rehydrated in HK buffer (20 mM HEPES (pH 7.5), 150 mM KCl) and subjected to a series of freeze–thaw cycles before extrusion through polycarbonate membranes (Whatman) with pore sizes of 100 or 400 nm using an Avanti Mini-Extruder. For GUV formation, lipid mixtures (DOPC:DPPC:cholesterol:Rhodamine-PE:Topfluor-cholesterol at 39:39:20:1:1) were dried on a ITO-coated glass slide and electroformation was conducted *via* applying 3.5V for 3 hour in 400 mM sucrose at 50 °C. To monitor the effect of tension changes on lipid phase separation, freshly prepared GUVs were subjected to isotonic buffer (400 mM glucose), hypertonic buffer (500 mM glucose) or hypotonic buffer (300 mM glucose) and imaged with inverted fluorescence microcopy immediately.

### Myotube stretching and imaging

C2C12 cells seeded on lamnin-coated BioFlex Culture Plate (Flexcell) was transfected with PABD-GFP and differentiated into myotube by DM incubation. After five days of differentiation, Rhodamine-Dextran was added into the medium to reach 1 µg/ml, and myotube was subjected to static, radial stretching with 20% strain by a Flexcell FX-5000T Tension System for 2 min and followed by no stretching for 2 mins; whereas control cells were incubated in Rh-Dextran containing medium for 4 mins without mechanical stress. Samples were then washed with PBS for 5 times, fixed with 4% formaldehyde for 30 min, cut from the plates and mounted with mounting medium and coverslip. PABD-GFP and Rh-Dextran signals were imaged with confocal microscopy (Zeiss, LSM700).

### Dynamin purification and activity analysis

Recombinant human Dyn2 was purified from insect cells and stored at −80 °C as previously described (Liu et al., 2011b). To determine membrane binding, 0.5 µM Dyn2 were incubated with 150 µM, 400 nm-sized liposomes with different amount of PA at 37 °C for 15 min. After centrifugation at 20,000 xg for 30 min, liposome-bound Dyn2 was pelleted down and was quantified by SDS-PAGE analysis. To analyze GTPase activity, GTP hydrolysis by Dyn2 was measured as a function of time using a colorimetric malachite green assay to detect the release of inorganic phosphate (Leonard et al., 2005). Briefly, 0.5 µM Dyn2 was incubated with 150 µM liposomes of different PA concentrations in a buffer containing 20 mM HEPES, pH 7.4, 150 mM KC, 1 mM GTP and 2 mM MgCl_2_. Aliquots were taken at several time points, free phosphate was determined using malachite green, and rates of hydrolysis were calculated. Fission activity of dynamin was measured as previously reported (Liu et al., 2011b). Briefly, the supported bilayers with excess membrane reservoir (SUPER) templates with different amount PA were incubated with 1 mM MgCl_2_, 1 mM GTP, and indicated dynamin for 30 min at room temperature. The released vesicles were separated from SUPER template by 260 ×*g* swing-out centrifugation. The fission activity is expressed as percentage of total fluorescence on SUPER templates.

### Transmission electron microscopy

To visualize the assembly of Dyn2 on liposomes, 1.0 μM dynamin was mixed with 25 μM liposomes with different lipid compositions and incubated at 37 °C for 10 min. The mixture was then adsorbed onto carbon-coated, glow discharged grids and stained with 1% uranyl acetate. Images were collected using a Hitachi H-7650 EM at 75 kV and a nominal magnification of × 120,000.

### Statistical analysis

Quantitative data were expressed as mean ± SD of at least three independent experiments. All data were analyzed with One-way ANOVA, except the DRM isolation result was analyzed with Student’s *t-*test. *p* < 0.05 was considered as statistical significance *p* values were indicated as *, *p* < 0.05; **, *p* < 0.01; ***, *p* < 0.001.

## Acknowledgements

We thank Dr. Do Sik Min (Pusan National University) for the GFP-PLD1 and GFP-PLD2 plasmids. We are grateful to Dr. Chau-Hwang Lee (Academia Sinica) for the assistance in GUV preparation and Dr. Damon Hoff (University of Michigan) for assistance with STORM imaging. We also thank the Single Molecule Analysis in Real-Time (SMART) Center of the University of Michigan, seeded by NSF MRI-R2-ID award DBI-0959823 to Nils G. Walter. This work was supported by Ministry of Science and Technology grant 107-3017-F-002-002 and National Taiwan University grant NTU-CDP-106R7808 to Y.W. Liu and NSF-1612917 to A.P. Liu.

## Author contributions

All authors participated in experimental design. JL, JJ, MCC, SSL, YCC, YAS and YWL performed experiments. JL, JJ, APL and YWL analyzed data and wrote the manuscript. APL and YWL supervised the project.

## SUPPLEMENTARY INFORMATION

**Inventory of supplementary information (SI):**

1. Three supplementary figures (Fig S1 is related to Fig1, Fig S2 is related to Fig 3, Fig S3 is related to Fig 4, Fig S4 is not related to any figure);

2. Three supplementary movies (these movies are related to Fig. 5).

**Figure S1.**
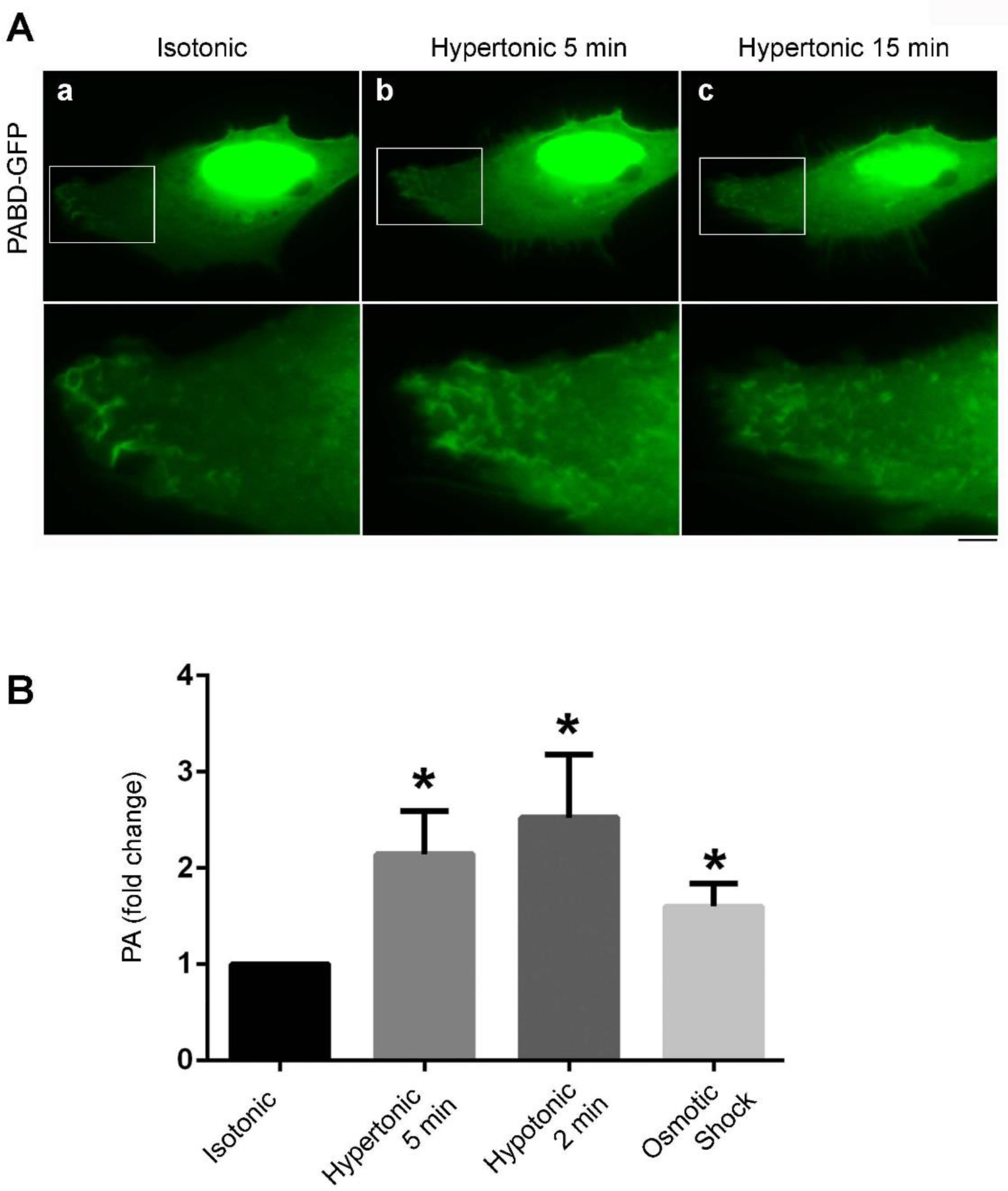
PA distribution and quantification upon tension manipulation. (A) Distribution of PA was monitored with PABD-GFP in myoblast treated with hypertonic buffer for 5 or 15 min. (B) Effects of different osmotic buffers on cellular PA amount. Myoblasts treated with indicated buffers were harvested and extracted for total lipids. Total PA/protein ratio were compared with the isotonic buffer-treated cells. *, *p* < 0.05. Scale bar, 10 µm.

**Figure S2.**
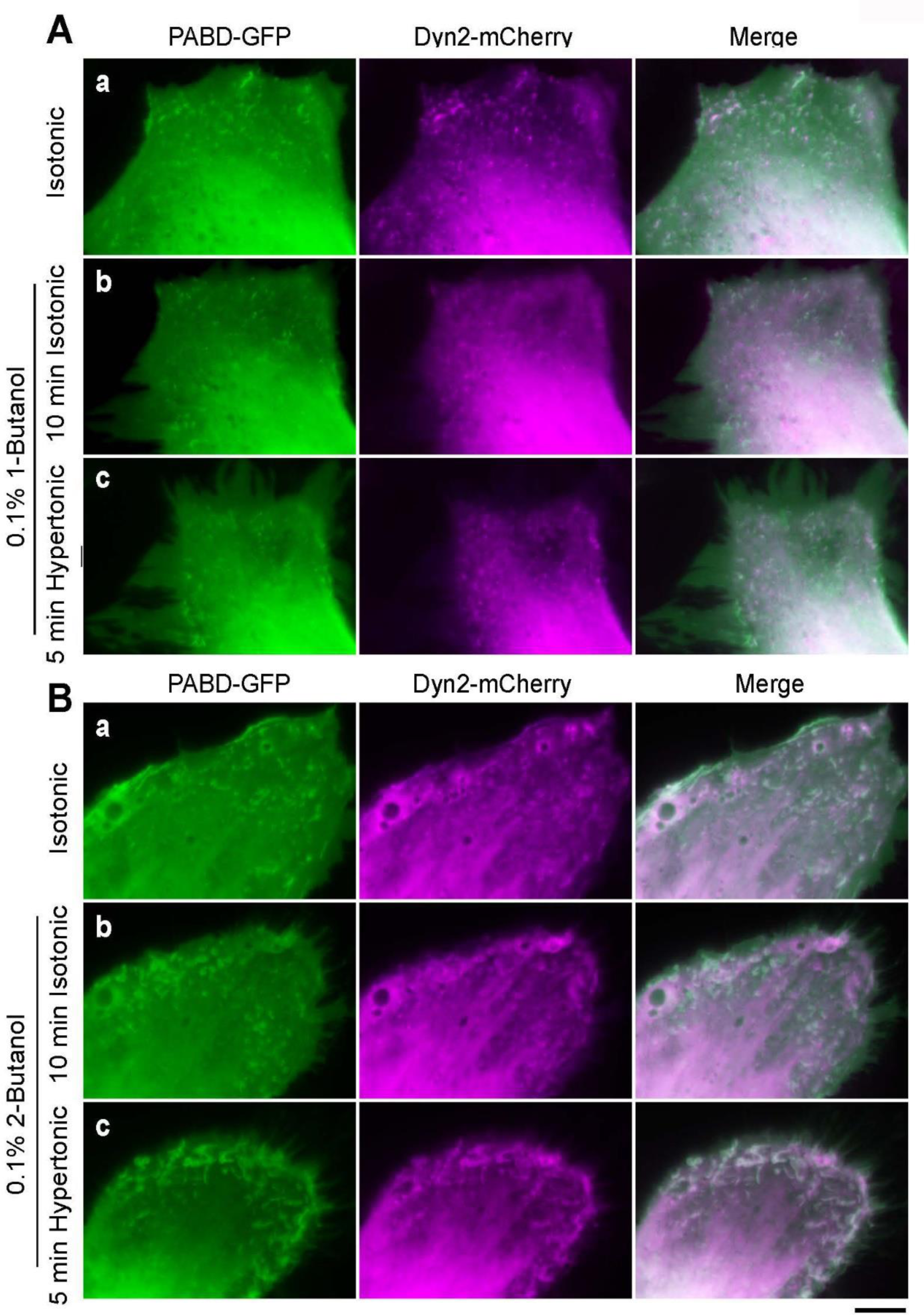
Effects of butanol on tension-induced, PA-rich membrane ruffling. PABD-GFP and Dyn2-mChery transfected myoblasts were pretreated with 0.1% 1-butanol (A) or 2-butanol (B) in isotonic buffer for 10 min and subjected to hypertonic buffer incubation in the presence of indicated alcohol. Scale bar, 10 µm.

**Figure S3.**
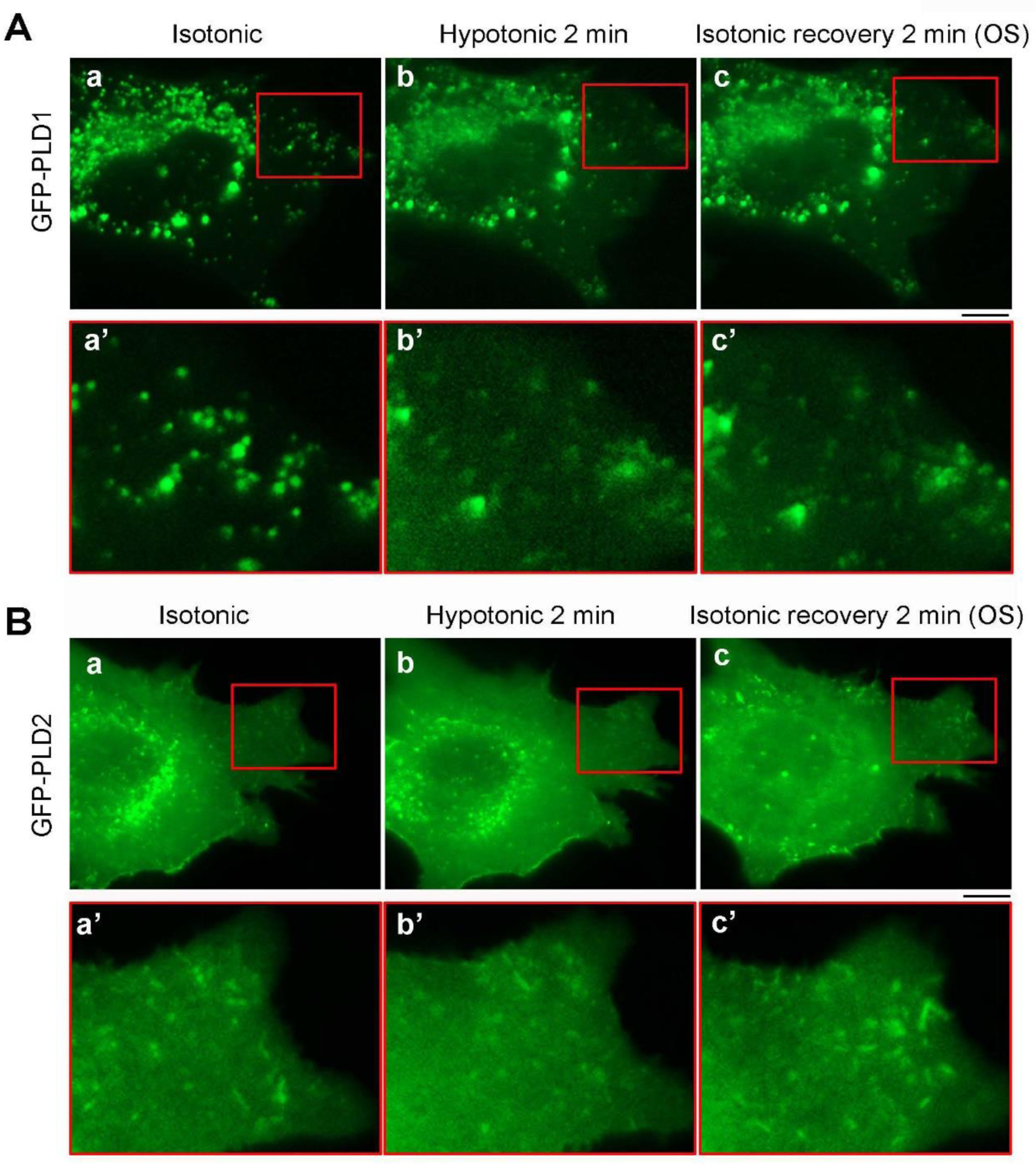
Subcellular distribution of GFP-PLD1 or GFP-PLD2 upon OS treatment. Myoblast expressing GFP-PLD1 or GFP-PLD2 were imaged in isotonic buffer (a), hypotonic buffer (b) and finally in isotonic recovery buffer (c) to monitor the kinetic distribution of these two PLDs. Magnified images of the boxed area are shown in the lower panel. Scale bar, 10 µm.

**Figure S4.**
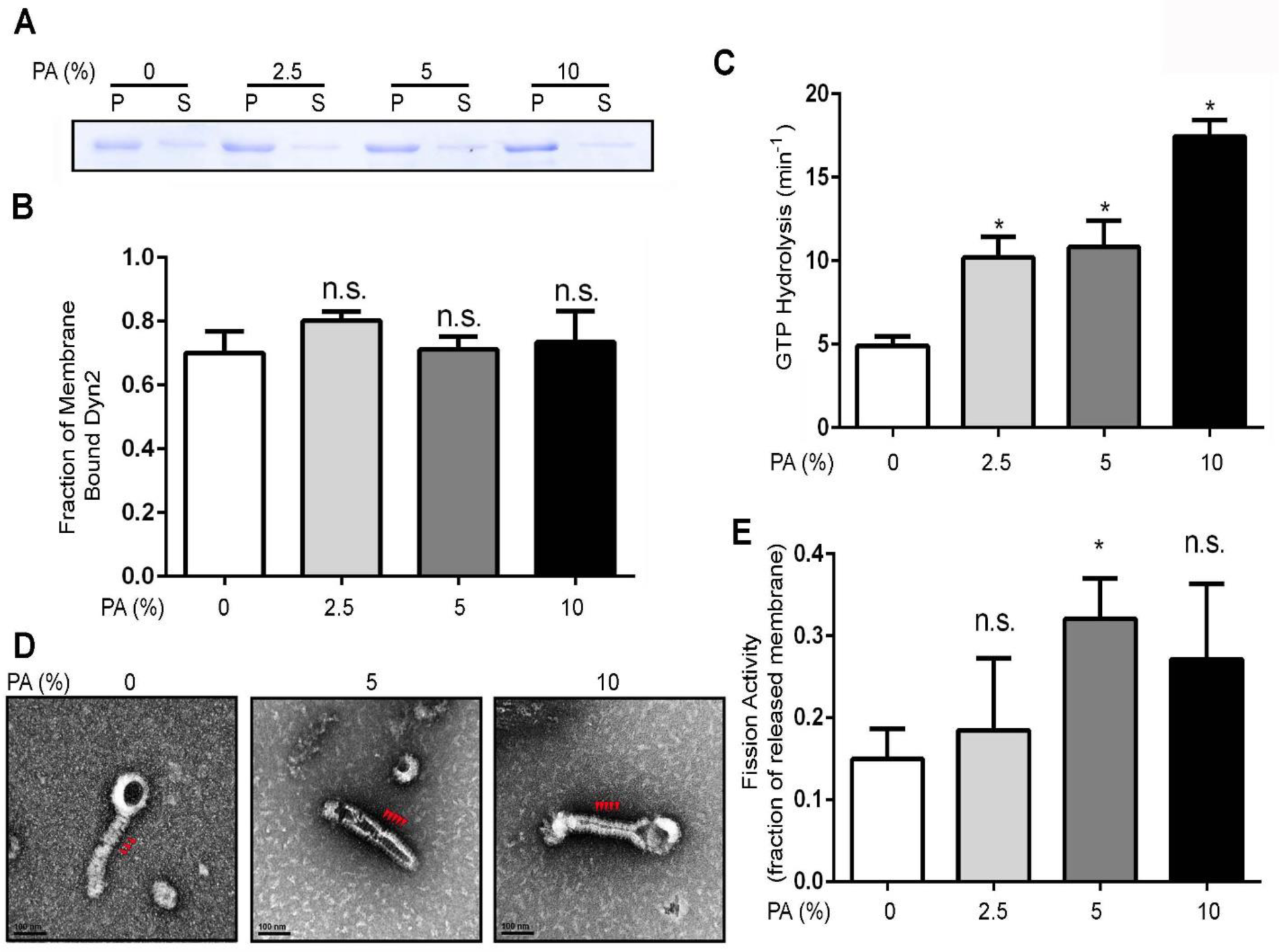
Effects of PA-containing membrane on Dyn2 activity. (A) Liposome binding assay. 0.5 μM purified human Dyn2 was incubated with 150 μM, 400 nm liposomes of indicated mole concentration of PA at 37°C for 15 min and sedimented into pellet (p) and supernatant (s) fractions by centrifugation. A representative Coomassie blue stained gel of p and s fractions of different percentage of PA-containing liposomes is shown. (B) The fraction of Dyn2 bound to liposomes were quantified with SDS-PAGE electrophoresis, Coomassie blue staining and ImageJ quantification. (C) Lipid-stimulated GTPase activity of Dyn2. 0.5 μM Dyn2 was incubated with 400-nm liposomes containing different PA concentration in the presence of 1 mM GTP at 37 °C. Released Pi was determined using a colorimetric malachite green assay. (D) Electron micrographs of Dyn2 assembled onto 100-nm liposomes with varying concentrations of PA. Red arrowheads indicate Dyn2 spirals. (E) Membrane fission activity of Dyn2. 0.5 μM Dyn2 was incubated with SUPER templates containing varying amounts of PA for 30 min in the presence of GTP. Membrane fission was measured by the release of fluorescently labeled vesicles into the supernatant after centrifugation of the SUPER templates. The data was normalized to its total lipids amount in the template. Data are shown as average ± SD (n=3). *, *p* < 0.05.

